# Targeted decontamination of sequencing data with CLEAN

**DOI:** 10.1101/2023.08.05.552089

**Authors:** Marie Lataretu, Sebastian Krautwurst, Matthew R. Huska, Mike Marquet, Adrian Viehweger, Sascha D. Braun, Christian Brandt, Martin Hölzer

## Abstract

Many biological and medical questions are answered based on the analysis of sequence data. However, we can find contamination, artificial spike-ins, and overrepresented rRNA sequences in various read collections and assemblies. In particular, spike-ins used as controls, as those known from Illumina or Nanopore data, are often not considered as contaminants and also not appropriately removed during analyses. Additionally, removing human host DNA may be necessary for data protection and ethical considerations to ensure that individuals cannot be identified.

We developed CLEAN, a pipeline to remove unwanted sequences from both long- and short-read sequencing techniques. While focusing on Illumina and Nanopore data with their technology-specific control sequences, the pipeline can also be used for host decontamination of metagenomic reads and assemblies, or the removal of rRNA from RNA-Seq data. The results are the purified sequences and sequences identified as contaminated with statistics summarized in a report.

The output can be used directly in subsequent analyses, resulting in faster computations and improved results. Although decontamination seems mundane, many contaminants are routinely overlooked, cleaned by steps that are not fully reproducible or difficult to trace. CLEAN facilitates reproducible, platform-independent data analysis in genomics and transcriptomics and is freely available at https://github.com/rki-mf1/clean under a BSD3 license.

**GRAPHICAL ABSTRACT:** 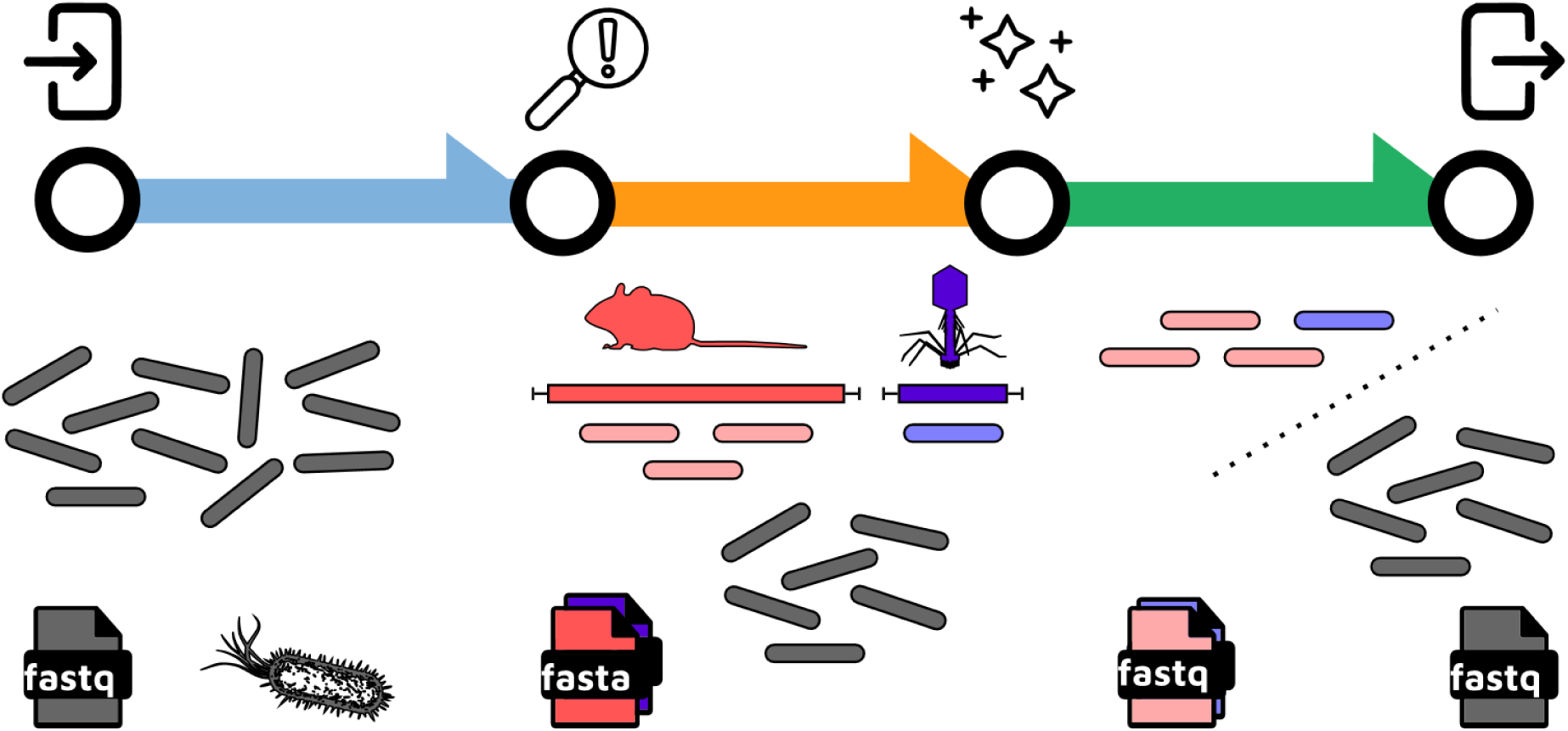

## INTRODUCTION

Next-Generation Sequencing (NGS) and Third-Generation Sequencing (TGS), commonly referred to as sequencing technologies, require quality control of the raw data. While most bioinformatic tools trim low-quality bases and remove adapter sequences, a critical step often overlooked is identifying DNA/RNA contamination from multiple sources [1], which can occur naturally or due to various factors. Contamination can arise during sample collection, shipping, or the sequencing library preparation process [2,3]. Additionally, control sequences are often introduced to calibrate basecalling and monitor run quality. However, these controls have been found to contaminate microbial isolate genomes in public databases: Illumina’s spike-in sequence, PhiX, were shown to be large-scale contaminants in microbial isolate genomes because the reads were not removed before assembly [4]. Also, we found that the positive control in Oxford Nanopore Technologies (ONT) DNA sequencing, a 3.6 kb standard amplicon known as DCS mapping the 3’ end of the Lambda phage genome, is mislabeled as *E. coli* or *Klebsiella quasipneumoniae* subsp. *similipneumoniae* plasmid in the NCBI GenBank (CP077071.1, CP092122.1), see **Supplementary Figures 1-3**. For ONT native RNA sequencing, a yeast ENO2 Enolase II transcript of strain S288C, YHR174W, functions as a positive control. Spike-in steps are usually optional; however, the information about whether a spike-in was used or not often does not reach the user working with raw reads.

Aside from addressing intentionally introduced control sequences and known contaminants, there are cases where specific biological sequences must be removed. For instance, in Illumina RNA-Seq samples, it’s often essential to remove ribosomal or mitochondrial RNA before read-count normalization and differential gene expression estimation [5]. This is particularly crucial for non-model species without optimized rRNA depletion kits [6]. Another example is eliminating host sequences, e.g. human sequences in human gut microbiome sequencing data [7].

Numerous tools have been developed for sequence data classification and decontamination, including Kraken 2 [8], Clark [9], Kaiju [10], HoCoRT [11] and Decontam [12], each with its own focus. Other tools specifically target ONT DNA spike-ins, such as nanolyse [13]. Nevertheless, despite the potential benefits in runtime and accuracy, many studies neglect proper read decontamination. As a direct result, we can find contamination omnipresent in genomic resources [14]. One reason might be that the output files of many pipelines cannot be directly used for downstream steps such as assembly or annotation and additional formatting of the files and extraction of the results are needed. We need decontamination tools that can be easily integrated into modern bioinformatics workflows.

To address these challenges, we introduce CLEAN (https://github.com/rki-mf1/clean), an all-in-one decontamination pipeline for short reads, long reads, and any FASTA-formatted sequences. Initially designed for removing Illumina and Nanopore spike-ins and host sequences in metagenomics, CLEAN’s functionality has been extended to also remove user-provided reference sequences. It also simplifies rRNA removal from Illumina RNA-Seq data and offers a streamlined QC report. CLEAN produces intermediate mapping files for further analysis and can be executed easily on various platforms. It uses common output formats, enabling direct integration into downstream analyses, enhancing decontamination in molecular biology research and genomic resources.

## MATERIAL AND METHODS

### Implementation

We use the workflow manager Nextflow v21.04.0 or higher [15] to manage our workflow, encapsulating each step in Docker [16], Singularity containers, or Conda environments [17] to ensure reproducibility of the results. The modular structure makes it easiler for us to update the containers and environments used by CLEAN periodically. CLEAN can be easily installed with a single command - the only prerequisites are Nextflow and one of Docker, Singularity, or Conda. We offer configurations for local execution, LSF and SLURM workload managers, and a simple cloud execution.

### Workflow

CLEAN’s input can be single- and paired-end Illumina FASTQ files, ONT or PacBio FASTQ read files, or FASTA files (**Figure 1**). The only required parameter is the input file, and users have the option to include a custom contamination reference FASTA file. We provide different external resources for common use cases, e.g. common host genomes, rRNA contamination reference and spike-in sequences (see **Supplementary Methods**). CLEAN combines all specified contaminants, allowing users to clean both host and spike-in reads in a single step. By default, each input file (FASTQ and/or FASTA) is mapped against the reference with minimap2 v2.26 [18] and specific options for short-read or long-read data. For Illumina, we also offer a kmer-based filtering option with bbduk (sourceforge.net/projects/bbmap) which directly results in clean and contaminated FASTQ files. Alternatively, the user can switch to BWA MEM [19] as short-read mapper. After mapping, we separate mapped from unmapped reads/contigs by the primary alignment with SAMtools [20]. CLEAN generates quality reports for input, clean and contamination files using FastQC (www.bioinformatics.babraham.ac.uk/projects/fastqc/) for Illumina, NanoPlot [21] for long reads, or QUAST [22] for FASTA files. MulitQC [23] summarizes all quality reports and mapping statistics in an HTML report. If minimap2 or BWA MEM were used, CLEAN additionally produces indexed mapping files (BAM) and an indexed contamination reference. If necessary, users can further analyze the results in a genome browser.

**Figure 1.**
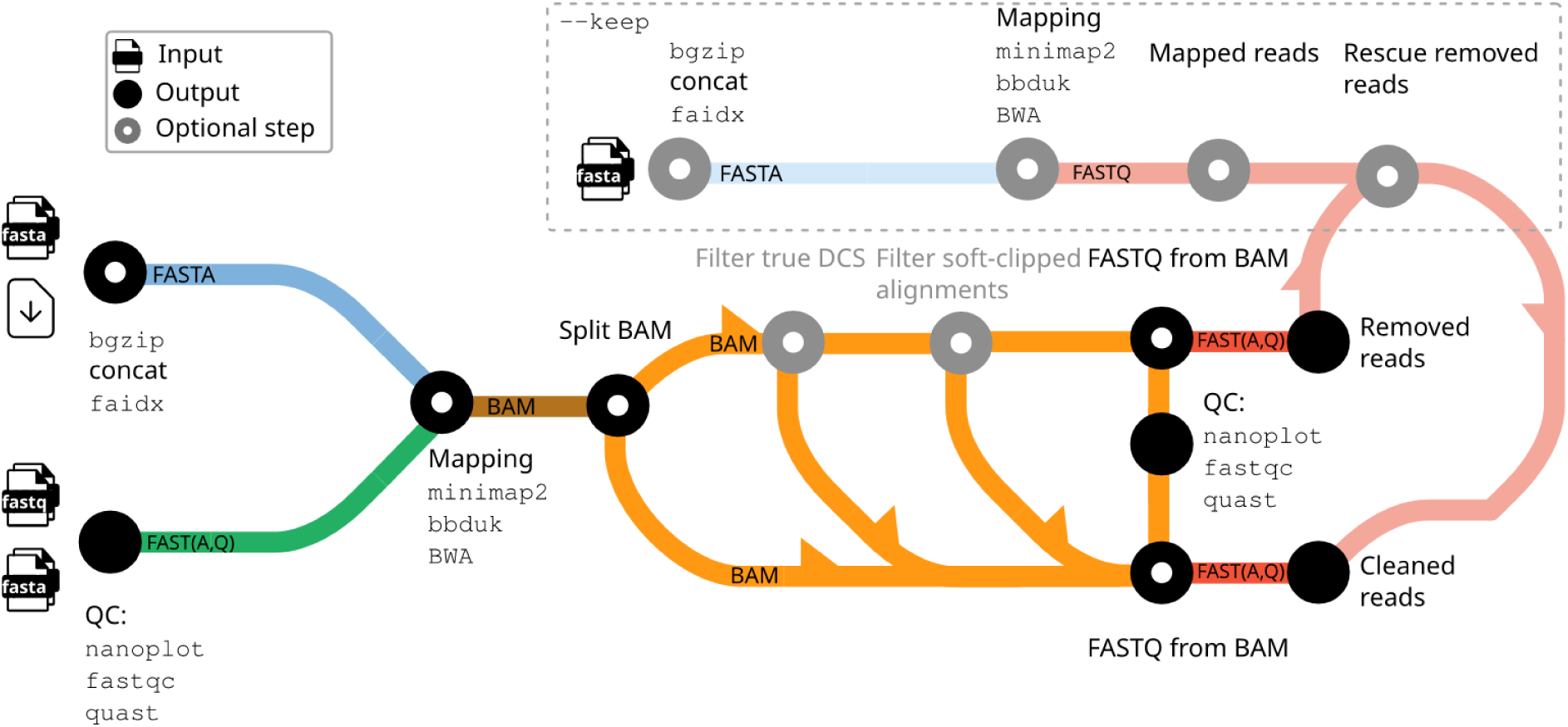
Schematic overview of the CLEAN workflow. Gray/blurred elements are optional and depend on the user input. The pipeline can search multiple FASTA or FASTQ inputs against a user-defined set of reference sequences (potential contamination). CLEAN automatically combines different user-defined FASTA reference sequences, built-in spike-in controls, and downloadable host species into one mapping index for decontamination. The user can also specify FASTA files comprising sequences that should explicitly not be counted as contamination. The output is finally filtered to provide well-formatted FASTA or FASTQ files for direct downstream analyses. The icons and diagram components that make up the schematic view were originally designed by James A. Fellow Yates & nf-core under a CCO license (public domain).

We want to highlight three parameter options: Firstly, for ONT data and DSC control, -- dcs_strict, which exclusively considers reads that align to the DCS and cover at least one of its artificial ends as contamination. This prevents inadvertent removal of similar phage DNA that might actually belong to a metagenomics sample. Secondly, with --min_clip, mapped reads are filtered by the total length (sum of both ends) of the soft-clipped positions. If --min_clip >= 1, the total number is considered, else the fraction of soft-clipped positions to the read length. Thirdly, the user can specify FASTA files with --keep. Input reads are then separately mapped to this reference. If a read maps to the “keep”-reference but was classified as contamination before, CLEAN moves the read to the set of clean reads. This feature helps mitigate false contaminant, particularly when dealing with closely related species or metagenomic samples.

### Test data sets and computations

We applied CLEAN to five distinct case studies to evaluate the pipeline’s ability to decontaminate sequencing data. Detailed descriptions of the underlying public sequencing data sets, novel sequencing data generated for this study, and all computational steps can be found in the **Supplementary Methods**.

#### Case study I: Removal of cell cultivation contamination

Nanopore and Illumina data from two previously published *Chlamydiifrater* isolates [24] and two novel isolates were decontaminated using a combined reference comprising the genome of *Chlorocebus sabaeus* (green monkey) and the mitochondrial DNA genome of *Chlorocebus pygerythrus* to remove host-derived contamination resulting from cultivation. Subsequently, we used the cleaned data to reconstruct hybrid *de novo* assemblies.

*Case study II: Decontamination in nanopore native RNA sequencing* Direct RNA sequencing data from HCoV-229E-infected Huh7 cells [25] were processed using CLEAN to distinguish viral, yeast, and human reads, with results compared to manual assignment.

#### Case study III: rRNA removal from Illumina RNA-Seq data

CLEAN’s performance for ribosomal RNA removal was assessed against SortMeRNA [26] using seven simulated Illumina datasets and one real RNA-Seq sample from a bat transcriptome study [6], with runtime and accuracy comparisons.

#### Case study IV: Human DNA spike-in removal from bacterial isolates after nanopore adaptive sequencing

Mixed DNA samples with varying human-to-bacteria ratios, spanning five bacterial species, were sequenced on the Nanopore platform using adaptive sequencing to deplete human sequences in real-time. Then, we used CLEAN to remove remaining human contamination after adaptive sequencing, facilitating accurate taxonomic classification and comparability between samples.

#### Case study V: Large-scale SARS-CoV-2 data decontamination

We used CLEAN to process 3866 SARS-CoV-2 nanopore amplicon data sets to ensure the removal of human contamination (including for personal data protection) while retaining viral reads using the “keep” parameter for subsequent upload to the European Nucleotide Archive. For scalability, the pipeline was executed on a high-performance computing cluster.

## RESULTS

### Case study I: Removal of cell cultivation contamination from Nanopore- and Illumina-sequenced Chlamydiaceae

The polished assemblies based on cleaned reads reveal 1.19 Mb circular genomes and 6 kb plasmids for each of the four *Chlamydiifrater* isolates. Without prior decontamination of the cell line DNA, contigs belonging to *Chlorocebus* species can be found in the final assemblies. Using an older version of Unicycler, running the assemblies without a CLEAN step of the raw read data also yields more fragmented final assembly results, likely due to the inflated complexity of the initial short-read graph. However, this issue was resolved by using a newer version of Unicycler, but still, contigs belonging to the host cell line could be found. Thus, decontamination of DNA belonging to a host cell line can improve the general assembly process and results in a much cleaner assembly.

### Case study II: Yeast enolase is a highly abundant spike-in control in Nanopore native RNA-Seq data

Nanopore sequencing is currently the only technology that allows the sequencing of native RNA strands without requiring a cDNA intermediate [27]. This ‘direct RNA’ protocol includes the addition of a calibration strand (amplified RNA sequences of the *S. cerevisiae* Enolase 2 mRNA, GenBank, NP_012044.1) as a spike-in positive control. Depending on the concentration of sample input RNA, this spike-in can represent a substantial fraction of the sequenced reads. In our study of direct RNA sequencing of Human Coronavirus genomes [25] these sequences made up 15.8% and 10.2% of the two samples, respectively. Due to algorithmic advances, re-basecalling the raw data with version 4.0.11 of the Guppy basecaller (RNA models are unchanged since then) yields more reads and a higher fraction of spike-in reads (31.4% and 31.0%, see **Figure 2**). Guppy does not filter these with default parameters but has an optional parameter (--calib_detect) to enable detection and filtering calibration strand reads. However, we found that this functionality does not adequately detect spike-in reads: 35.4% and 19.8% of spike-in reads were still present when using this parameter. Applying CLEAN to this dataset removes all calibration strand reads (see **Figure 2**).

**Figure 2.**
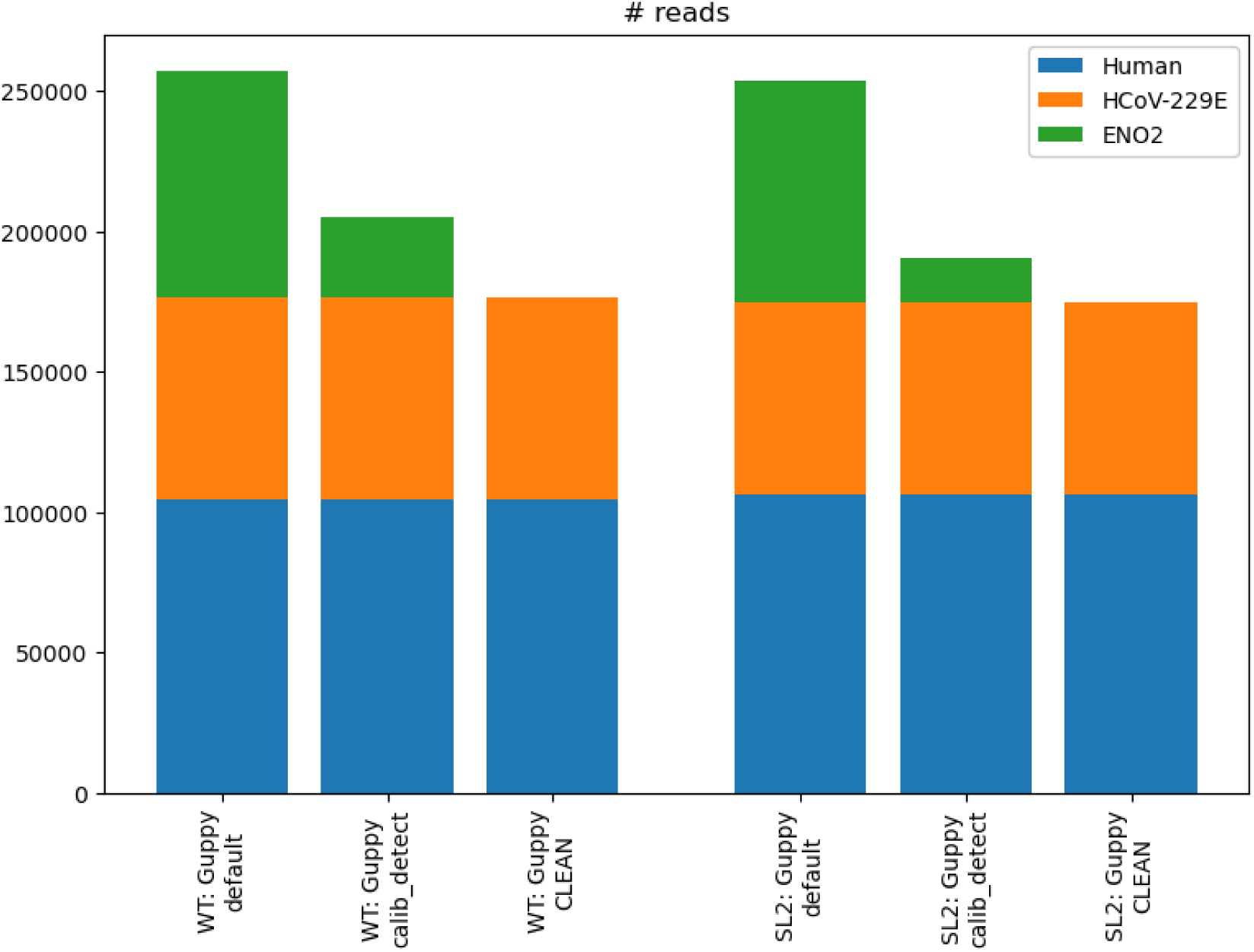
Number of reads mapping to the human genome, HCoV-229E or *S. cerevisiae* Enolase 2 (from bottom to top) for two HCoV-229E samples WT (left) and SL2 (right) after Guppy (default parameters), Guppy with --calib_detect or after CLEAN usage. Only CLEAN is able to remove all reads deriving from the dRNA control sequence. WT — wild type sample, SL2 — sample with different RNA secondary structure.

Generally, if a positive control is not needed for the experiment, we suggest skipping the addition of this spike-in. This can increase the yield of desired RNA reads by freeing up throughput capacity. For all direct RNA read data with added spike-in, we propose using CLEAN to remove these sequences reliably and quickly before downstream analyses are performed.

### Case study III: Speeding up an everyday task in transcriptomics – removal of rRNA from Illumina RNA-Seq data

CLEAN performs equally well compared to SortMeRNA for the non-rRNA datase: CLEAN’s recall is slightly better (< 0.001) than SortMeRNA’s. CLEAN’s recall for the six rRNA datasets is, on average, 0.03 lower (minimum 0.01, maximum 0.06) than SortMeRNA, see **Supplementary Table 1**. On the real-data sample, CLEAN runs about 1.7-fold faster than SortMeRNA, see **Supplementary Figure 4**. Results vary slightly with <0.014 % divergence.

### Case study IV: Decontamination of human spike-in DNA from bacteria isolates

Nanopore sequencing allows for the selective depletion or enrichment of target sequences in real time during the sequencing run. Here, we created an ONT dataset that includes five different bacterial species, to which four different concentrations of human DNA were added and sequenced with real-time depletion enabled. However, selective sequencing does not remove 100 % of the target (human) DNA, and in particular, results in shorter reads of the targeted sequence (as sequencing stops as soon as a classification is possible). Therefore, we used CLEAN to remove the remaining human DNA contamination from the bacterial samples.

Overall, we observed a 99.07 % reduction in human reads after applying CLEAN to the sequencing results. The resulting cleaned read data were classified with Kraken 2 (a k-mer-based method), to identify and quantify remaining human reads. **Supplementary Table 2** lists the total number of reads per sample, those that could be taxonomically classified as human using Kraken 2, and the percentage of reads remaining after applying CLEAN. To further validate the remaining ∼1 % human reads not removed by CLEAN (**Supplementary Figure 5**), we mapped them back to a reference human genome. Remarkably, only one-tenth could be successfully aligned to the genome, explaining why CLEAN’s mapping-based approach did not identify them as human. Therefore, we assume that the discrepancy might be related to the k-mer-based classification of Kraken 2 on these few remaining sequencing reads. **Figure 3** illustrates the amount of human DNA spike-in remaining in the *Acinetobacter pittii* (Ap) sample that could not be completely removed by selective sequencing and the level of decontamination achieved by CLEAN (see **Supplementary Figure 5** for all samples). Our results show that CLEAN can be used to effectively remove human contamination from bacterial datasets. However, a small number of reads can still be annotated as human, which could be due to algorithmic and/or reference biases.

**Figure 3.**
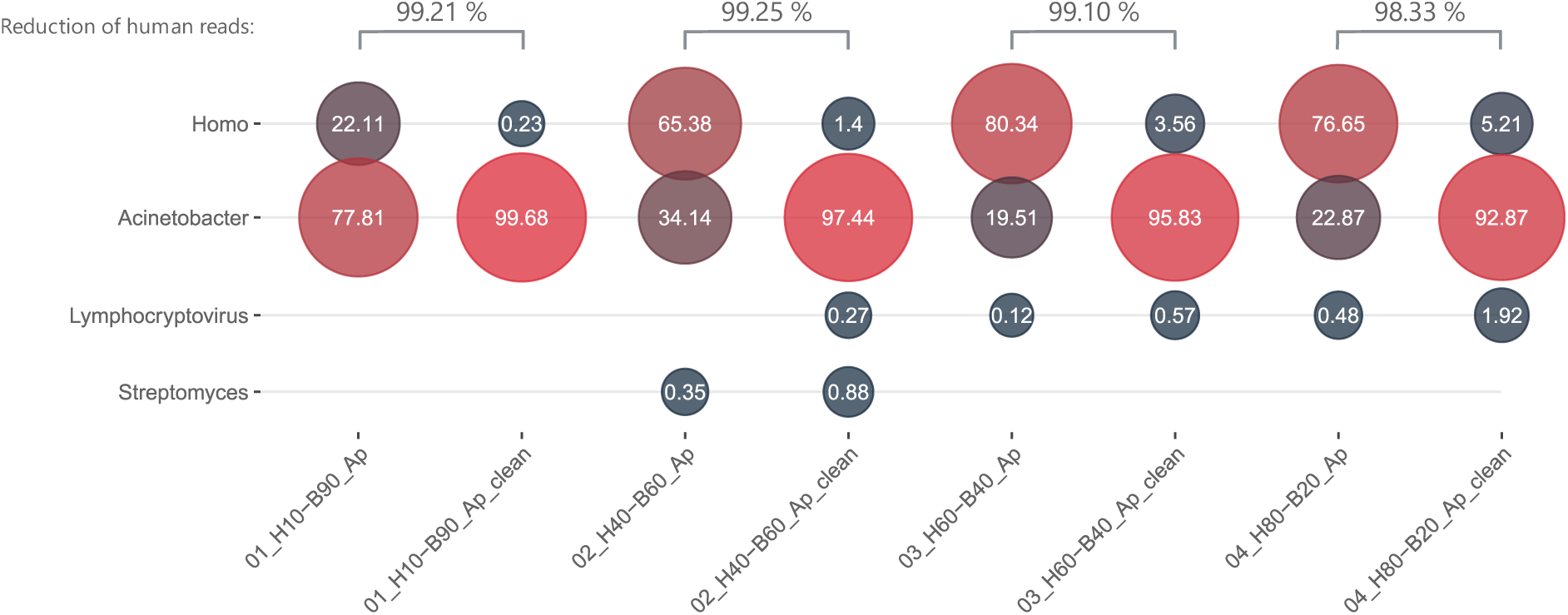
Abundances of reads per sequencing run before and after CLEAN classified with Kraken 2 and Bracken for *Acientobacter pittii* (Ap). H — percent of human DNA spike-in; B — percent of bacteria DNA. **Supplementary Figure 5** shows the results for all investigated bacteria.

### Case study V: Removal of human contamination from a large SARS-CoV-2 genomic surveillance data set

CLEAN processed all 3,866 SARS-CoV-2 amplicon FASTQ files on a high-performance cluster (HPC) in 2h 14min, having used 1,547 CPU hours for computation. Across all samples the mean number of reads that were removed was 274, and median number of reads removed was 61. Only 15 samples had more than 5 % of their reads removed.

Looking at reads that mapped to both the human genome as well as the SARS-CoV-2 genome, in all cases the reads mapped better to the SARS-CoV-2 genome: SARS-CoV-2 alignments were typically the full length of the read and with high similarity, while alignments to the human genome were only covering a small fraction of the reads. Because of this we did not consider those reads to be contamination and kept them using CLEAN’s --keep functionality.

## CONCLUSION

We developed CLEAN to easily screen any nucleotide sequences against reference sequences to identify and remove potential contamination. This includes common tasks such as the removal of positive controls added during library preparation, host contamination, or ribosomal RNAs. Decontamination with CLEAN can be easily used as a preprocessing step before a main analysis since the output needs no further processing or reformatting. By default, the pipeline uses alignment-based approaches for short- and long-reads that subsequently also allow for the inspection of the reads aligned to a potential contamination reference in more detail. Furthermore, CLEAN provides quality control reports for more insights. CLEAN is freely available at https://github.com/rki-mf1/clean and can be easily installed and executed using Nextflow.

### Limitations

CLEAN cannot be used to remove unexpected contaminants. For such a task, DecontaMiner, a tool to remove contaminating sequences of unmapped reads [28], or QC-Blind, a tool for quality control and contamination screening without a reference genome [29] can be used. Other tools try to find unexpected compositions in metagenomics samples to identify contaminants [12]. With CLEAN we did also not focus on the detection of cross-contamination. For this task other tools such as ART-DeCo [30] can be used. Furthermore, CLEAN should not be used in situations where specialized tools are available, e.g., SortMeRNA for rRNA annotation and Kraken 2 for taxonomic classification.

## DATA AVAILABILITY

CLEAN, including the user manual, is available on GitHub (https://github.com/rki-mf1/clean) under the open source BSD3 license. All supporting analysis scripts are available in OSF (doi.org/10.17605/OSF.IO/CUXEM). Data used in this work are available in public databases:

Case study I: SRA BioSample IDs: SAMEA6565319 (strain 15-2067_O50), SAMEA6565320 (strain 15-2067_O99) and ENA Study Accession ID: PRJEB59173 (strains 15-2067_O09 and 15-2067_O77)

Case study II: Sequencing data and scripts: doi.org/10.17605/OSF.IO/UP7B4 Case study III: SRA BioSample ID: SAMN10246232

Case study IV: ENA Study Accession ID: PRJNA1199779 Case study V: ENA Study Accession ID: PRJEB76939.

## SUPPLEMENTARY DATA

Supplementary Data are available online.

## AUTHOR CONTRIBUTIONS

MHö provided conceptualization, initial design, and a first implementation. ML optimized the pipeline code, realized the final implementation, conducted the experiments, and created figures. SK performed the benchmark for the Coronavirus dRNA-Seq experiment and provided corresponding results and methods. MHu performed the benchmark on SARS-CoV-2 samples and extended the functionality of the pipeline by extensive testing, bug fixing, and reorganization of the output structure. SB prepared bacteria samples with human spike-in DNA and performed nanopore sequencing; CB and MM analyzed the bacteria data set with CLEAN and provided methods and results. CB and AV provided the initial backbone code structure for the workflow. MHö and ML wrote the first draft of the manuscript. All authors actively participated in the writing and final editing of the manuscript. All authors have read and agreed to the published version of the manuscript.

## ACKNOWLEDGEMENTS

We thank Fabien Vorimore from ANSES, France for sequencing of the two *Chlamydiifrater* strains and providing the raw data for our benchmark. We thank Stephan Fuchs from RKI, Germany, for fruitful discussions.

## SUPPLEMENTARY METHODS

### Test data sets and computations

#### Case study I: Removal of cell cultivation contamination from Nanopore- and Illumina-sequenced Chlamydiaceae

We obtained Nanopore (FAST5) and Illumina (FASTQ) data of two recently defined *Chlamydiifrater volucris* isolates, 15-2067_O50 (SAMEA6565319) and 15-2067_O99 (SAMEA6565320) [1] and re-basecalled the Nanopore raw signal data with Guppy (v6.0.0 and SUP accuracy model). In addition, we obtained new data for two isolates (15-2067_O09 and 15-2067_O77), probably also belonging to the species *Chlamydiifrater volucris*, which were cultivated on a cell line derived from *Chlorocebus sabaeus* (Green monkey). DNA was extracted and sequenced with Oxford Nanopore and Illumina by colleagues at ANSES, France (Fabien Vorimore) and as described for the already published *Chlamydiifrater* strains [1]. We used CLEAN to decontaminate all reads against DCS (-- control dcs, for Nanopore) and phix (--control phix, for Illumina), *Chlorocebus sabaeus* (--host csa) and the mitochondrial genome of *Chlorocebus pygerythrus* (--own NC_009747.1). Unfortunately, it is not known which species of *Chlorocebus* was exactly used for the construction of this cell line [1]. Thus, we decided to use the complete chromosomal and mitochondrial genome of *C. sabaeus* and add the mtDNA of *C. pygerythrus* (no chromosomal sequences are available) to increase our chances for proper decontamination (CLEAN seamlessly allows the usage of multiple references). During our analyses, we also discovered that the mitochondrial DNA of *C. pygerythrus* provides an even better matching than the mtDNA of *C. sabaeus*. After decontamination, we length-filtered the ONT reads with filtlong (v0.2.0, parameters: --target_bases 1.2 * 200000000) (https://github.com/rrwick/Filtlong) and quality-trimmed Illumina reads with fastp (v0.20.1, parameters: -5 -3 -W 4 -M 20 -l 15 -x -n 5 -z 6) [2]. Finally, we *de novo* assembled the cleaned and filtered short- and long-reads with Unicycler (v0.5.0, default parameters) [3] followed by independently mapping the Illumina short reads with BWA (v0.7.17) [4] to the respective resulting Unicycler assembly and subsequent polishing the assembly with polypolish (v2.2.0) [5].

#### Case study II: Coronavirus native RNA sequencing with Nanopore

Virus generation, RNA isolation, sample preparation, and sequencing are detailed in [6]. Briefly, Huh7 cells were infected with recombinant HCoV-229E variants, yielding two samples in cell culture (WT (wild type) and SL2 (mutant)). Total RNA of these was isolated, and 1 µg of RNA in 9 µL was carried into the library preparation with the Oxford Nanopore direct RNA-Seq (DRS) protocol (SQK-RNA001). Sequencing ran for 48h on an R9.4 flow cell on a MinION device.

For this study, the raw data was basecalled with Guppy (version 4.0.11), once with and once without the --calib_detect parameter. Assignment of reads to either HCoV-229E, *S. cerevisiae* Enolase 2, or human was done by mapping to a combined reference of all three with minimap2 (version 2.17, parameters: -ax splice -k14).

Finally, we used CLEAN on the basecalled DRS reads with calibration strand detection and compared the results to the manual assignment. All commands and the plotting script are available at https://doi.org/10.17605/OSF.IO/CUXEM.

#### Case study III: rRNA removal from bulk RNA-Seq Illumina data

We tested and compared CLEAN’s functionality to remove ribosomal RNA against SortMeRNA (v4.3.4) [7]. All seven simulated datasets were downloaded from [7]. Briefly, 1 million single-end rRNA Illumina reads with a read length of 100 bp were simulated with different identities with respect to the SILVA database, or origin from truncated sections of the bacteria phylogenetic tree. One of the seven simulated samples contains non-rRNA reads. We converted the provided FASTA files into FASTQ files with seqtk (v1.3-r106, https://github.com/lh3/seqtk) and calculated the recall for each dataset considering rRNA reads as positive label for the six rRNA datasets and non-rRNA reads as positive label for the non-RNA dataset.

To compare runtime performance, we chose a non-simulated Illumina RNA-Seq sample (GEO Accession GSM3431091) from a bat transcriptome study [8]. For total RNA obtained from a bat (*Myotis daubentonii*) cell line, cDNA libraries were prepared utilizing the Illumina Ribo-Zero rRNA Removal Kit for human/mouse/rat. We used CLEAN with the --rrna parameter and SortMeRNA to remove rRNA reads from the sample.

We run each tool ten times with 30 threads to compare runtime differences measured with Linux’s time command on a Linux server (CPU: Opteron 6376, 64 x 2,1 GHz, RAM: 768 GB).

#### Case study IV: Decontamination of human spike-in DNA from bacteria isolates

Bacteria used in this study included *Staphylococcus aureus* (Sa), *Enterococcus faecium* (Ef), *Klebsiella pneumoniae* (Kp), *Acinetobacter pittii* (Ap), and *Escherichia coli* (Ec). Bacterial isolates were cultured overnight at 37 °C on Columbia blood agar. DNA was purified using Qiagen DNeasy Blood and Tissue Kit (Qiagen, Hilden, Germany) following the manufacturer’s instructions. Human DNA was obtained from the Genome in a Bottle (GIAB) project, specifically from the Ashkenazim Son sample (HG002/NA24385/huAA53E0), with accession number GCA_001542345.1, and was purchased from Sigma-Aldrich (order number: NIST8391, Taufkirchen, Germany). Bacterial DNA was mixed with human DNA in varying ratios of 10:90, 40:60, 60:40, and 80:20 (human/bacteria) to create artificial mixed samples. The Oxford Nanopore Technologies (ONT) MinION platform was used to sequence all 20 (5 bacteria x 4 human spike-in ratios) artificial samples containing bacterial and human DNA. Library preparation was performed using the 1D genomic DNA ligation kit (SQK-NBD114.24; ONT, Oxford, UK) following the manufacturer’s instructions for flow cell FLO- MIN114 containing R10.4.1 pores. Prior to library preparation, size selection was performed using AMPure beads (Beckman Coulter, Krefeld, Germany) in a 1:1 (v/v) ratio with all DNA samples. The flow cell was loaded with approximately 600 ng/µL of DNA (according to the Qubit 4 Fluorometer; Thermo Fisher Scientific, Waltham, MA, USA). Sequencing was conducted for 90 hours using MinKNOW software version 24.02.16, starting with a total of 1038 active pores. Live basecalling was performed using the dna_r10.4.1_e8.2_400bps_fast model, with live demultiplexing enabled during sequencing. Resulting FASTQ files were created every ten minutes and saved in folders named by the used barcodes. The depletion flag was set in the MinKNOW software to selectively deplete human DNA during sequencing and based on a database containing a human reference genome (accession GCF_000001405.40), reducing read lengths of human reads to approximate 400 bp. Sequenced raw data was taxonomically classified with Kraken 2 (v2.1.1) followed by Bracken (v2.8) using the standard RefSeq Database (https://benlangmead.github.io/aws-indexes/k2, October 2023).

#### Case study V: Removal of human contamination from a large SARS-CoV-2 genomic surveillance data set

We used CLEAN to prepare a large collection of 3866 SARS-CoV-2 virus Nanopore sequencing datasets to be made public on the European Nucleotide Archive (ENA). All samples were generated using amplicon sequencing, so the risk of contamination is lower than some other sample preparation methods. Nevertheless, it is important to ensure that no personally identifiable human data is present in datasets that are made public. This is a requirement both of ENA and also our own institute, to protect employees who may have inadvertently contaminated the samples. To achieve this goal we used CLEAN, setting the human reference genome (hg38) as the host. For this dataset, we also explicitly specified the SARS-CoV-2 reference genome (NCBI accession NC_045512.2) as the target genome we were trying to sequence using CLEAN’s --keep argument, so that CLEAN would not remove any reads that map well to both the SARS-CoV-2 and human genomes. We then did an analysis on these reads that map to both genomes to compare mapping quality and decide whether those reads would need to be removed.

CLEAN was run on all 3866 FASTQ files on a high-performance computing cluster using the SLURM cluster management system. The run was allowed to use up to 1000 parallel jobs spread across several compute nodes, each of which contains two AMD EMYC 7742 64-core processors and at least 1TB of RAM.

### External resources

The user can define a contamination reference or choose from included ones. These are the currently provided host genomes in CLEAN version v1.0.0-alpha: *Homo sapiens* (Ensembl release 99), *Mus musculus* (Ensembl release 99), *Gallus gallus* (Ensembl release 99), *Escherichia coli* (Ensembl release 45), *Chlorocebus sabaeus* (NCBI GCF_000409795.2), and *Columba livia* (NCBI GCF_000337935.1). A genome is only downloaded once on-demand and can be reused. The list of automatically downloadable references can be easily extended upon request or by experienced users. However, the user can also always provide additional reference FASTAs via a parameter. As an rRNA reference, we provide the rRNA database from SortMeRNA, a tool commonly used to filter rRNA from metatranscriptomic data. The database contains representative rRNA sequences from the Rfam and SILVA databases (see https://github.com/biocore/sortmerna/blob/master/data/rRNA_databases/README.txt). Spike-in sequences for direct RNA and DNA ONT sequencing are taken from Guppy, the basecaller developed by ONT: yeast enolase ENO2/YHR174W of 1.2 kb and a Lambda Phage amplicon of 3.6 kb. By further investigating the latter, we found another resource at the ONT community for the DCS sequence (https://assets.ctfassets.net/hkzaxo8a05x5/2IX56YmF5ug0kAQYoAg2Uk/159523e326b1b791e3b842c4791420a6/DNA_CS.txt). This 3560 nt long sequence is a substring of the Guppy sequence (3587 nt), where the first 27 nucleotides are duplicated at the start, see Supplemental Figure 2 and 3. The first 65 nt (Guppy 92 nt) and the last 48 nt seem to be artificial as they show no hits in a BLAST search against the NCBI nucleotide collection (nr/nt).

---

**Supplementary Figure 1.**
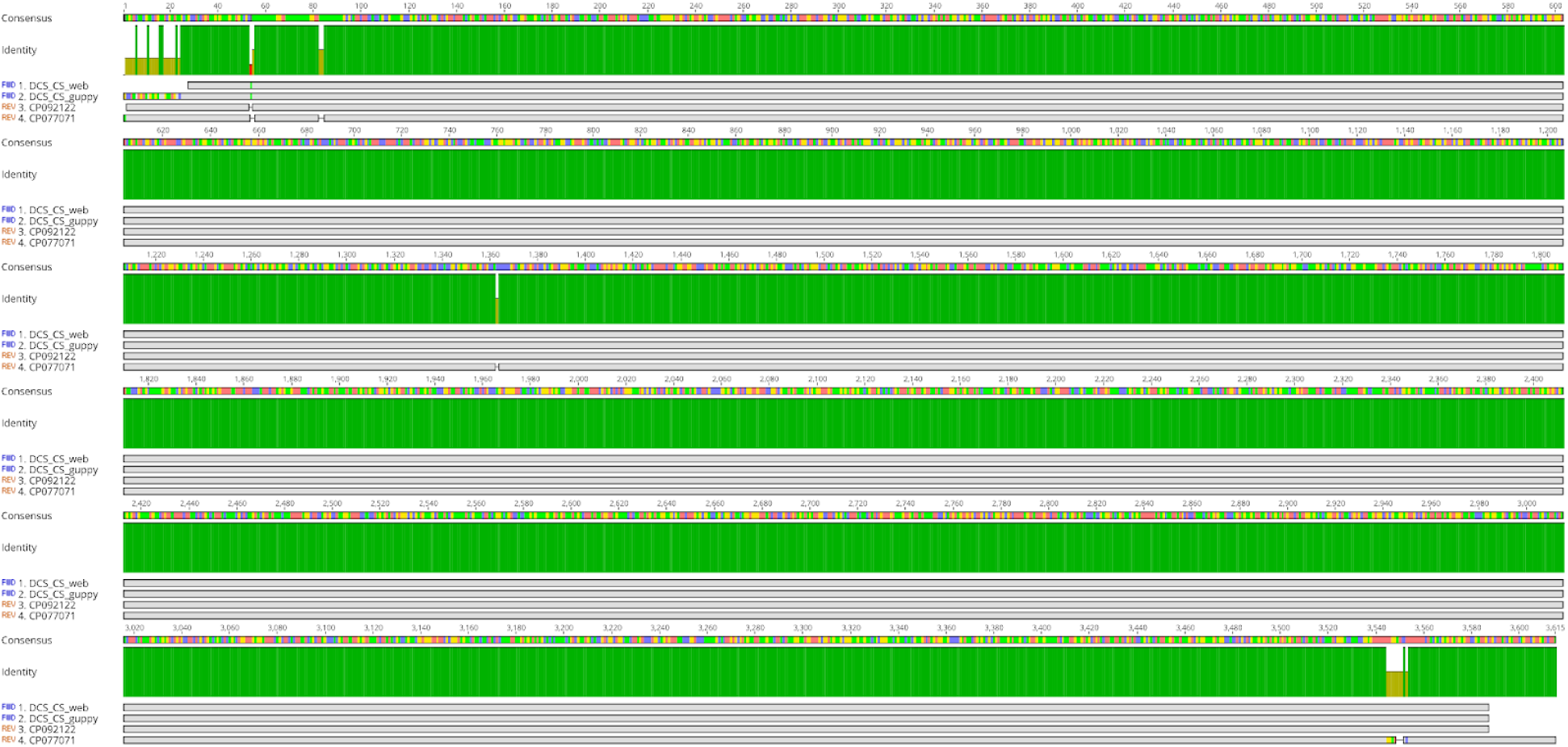
Geneious Prime (v2021.2.2, Geneious alignment, default parameters, https://www.geneious.com) alignment of *E. coli* (CP077071.1), *Klebsiella quasipneumoniae subsp. similipneumoniae* plasmids (CP092122.1) and DCS control sequences from Guppy (DCS_CS_guppy) and the ONT community (DCS_CS_web, https://assets.ctfassets.net/hkzaxo8a05x5/2IX56YmF5ug0kAQYoAg2Uk/159523e326b1b791e3b842c4791420a6/DNA_CS.txt). The high similarity suggests that both plasmids are contamination and falsely classified as plasmids.

**Supplementary Figure 2.**
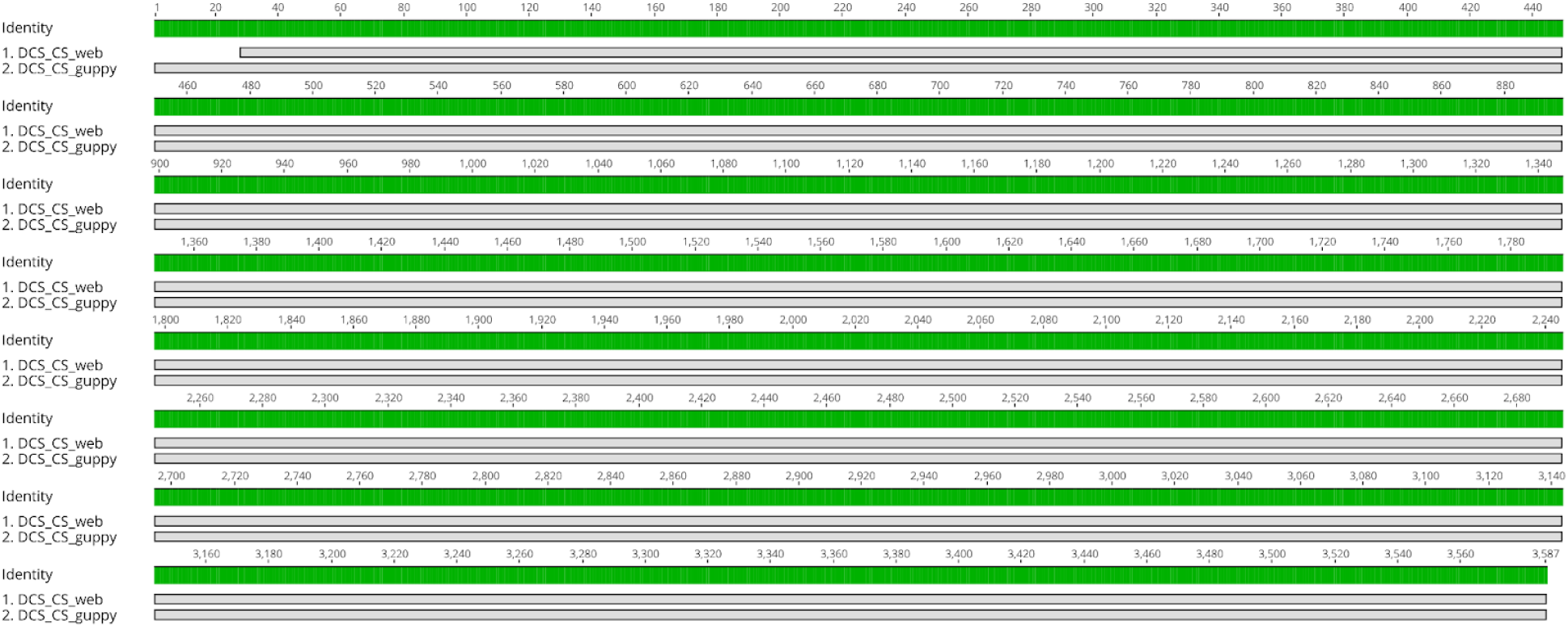
Geneious Prime (v2021.2.2, Geneious alignment, default parameters, https://www.geneious.com) alignment of DCS control sequences from Guppy (DCS_CS_guppy) and the ONT community (DCS_CS_web, https://assets.ctfassets.net/hkzaxo8a05x5/2IX56YmF5ug0kAQYoAg2Uk/159523e326b1b791e3b842c4791420a6/DNA_CS.txt). Sequences are identical except for the first 27 nt in the Guppy version, which are duplicated subsequently.

**Supplementary Figure 3.**
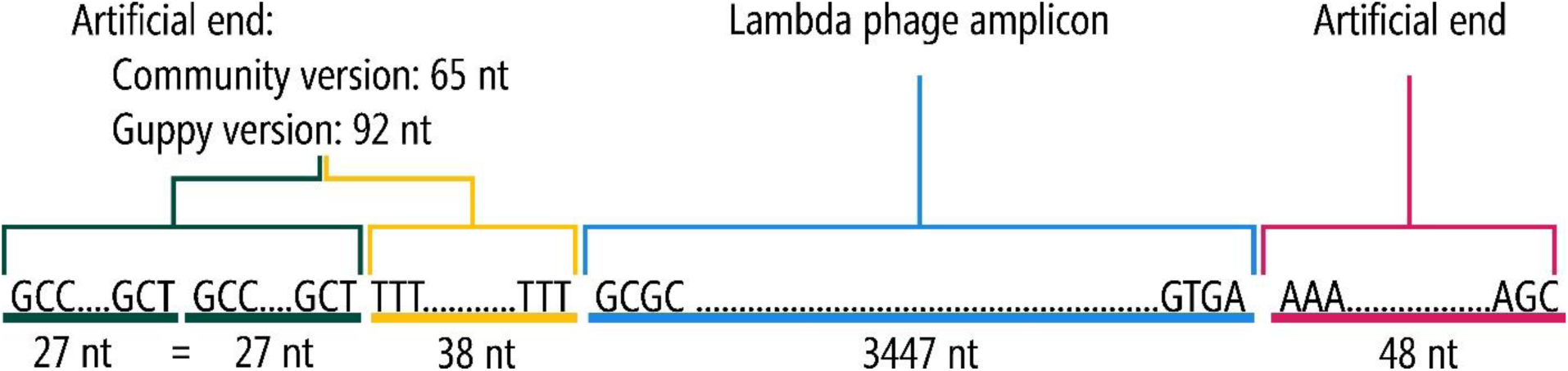
Schematic illustration of the DCS control sequence: artificial ends frame a part of the Lambda phage genome. Available sequences (ONT community and Guppy installation) differ by a duplication of the first 27 nucleotides.

**Supplementary Figure 4.**
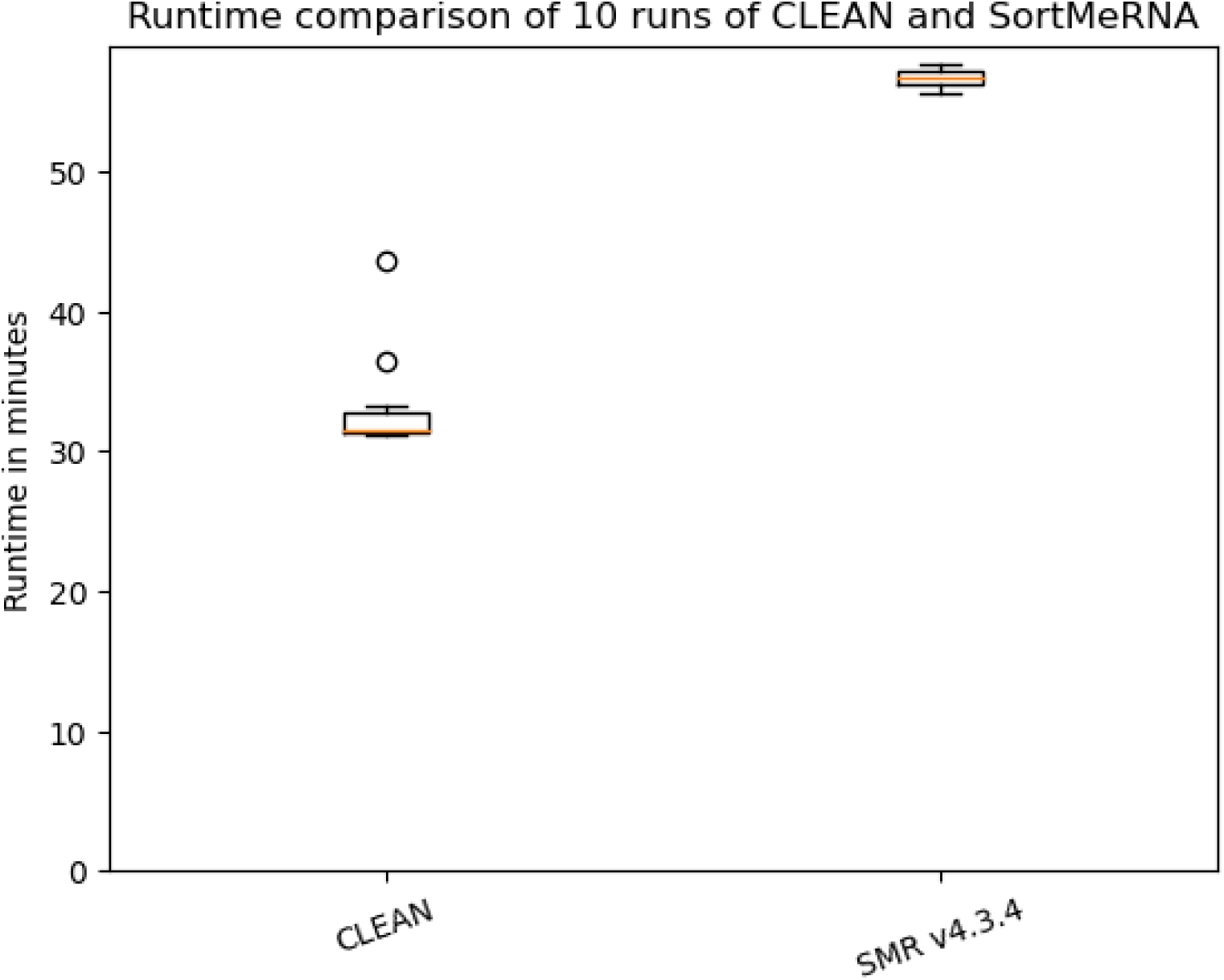
Comparison of the runtime for ten repeated runs of CLEAN and SortMeRNA v4.3.4. Both tools were executed on a Linux server (CPU: Opteron 6376, 64 x 2,1 GHz, RAM: 768 GB) with 30 threads. Time was measured with the Linux time command.

**Supplementary Figure 5.**
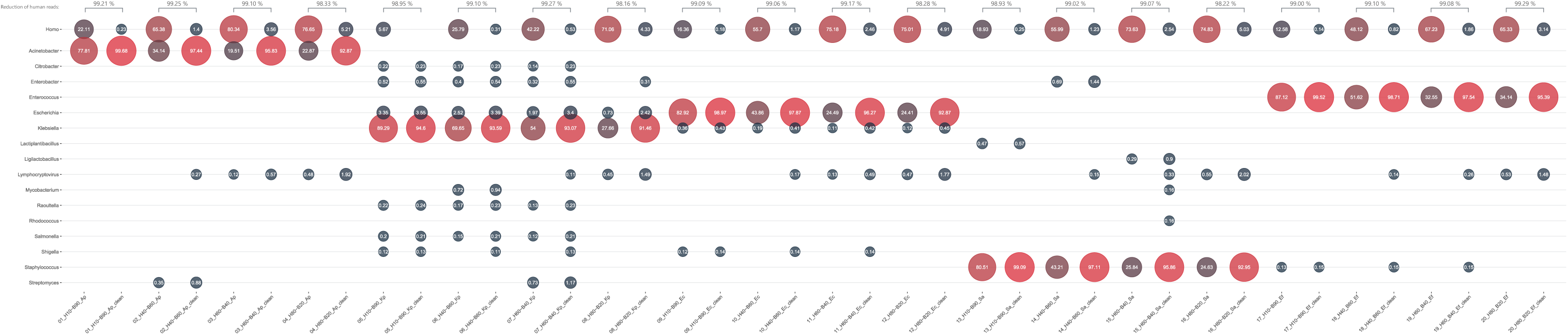
Abundances of reads per sequencing run before and after CLEAN classified with Kraken 2 and Bracken. H — % of human DNA spike-in; B — % of bacteria DNA; Ap — *Acientobacter pittii*; Kp — *Klebsiella pneumoniae*; Ec — *Escherichia coli*; Sa — *Staphylococcus aureus*; Ef — *Enterococcus faecium*.

**Supplementary Table 1.**
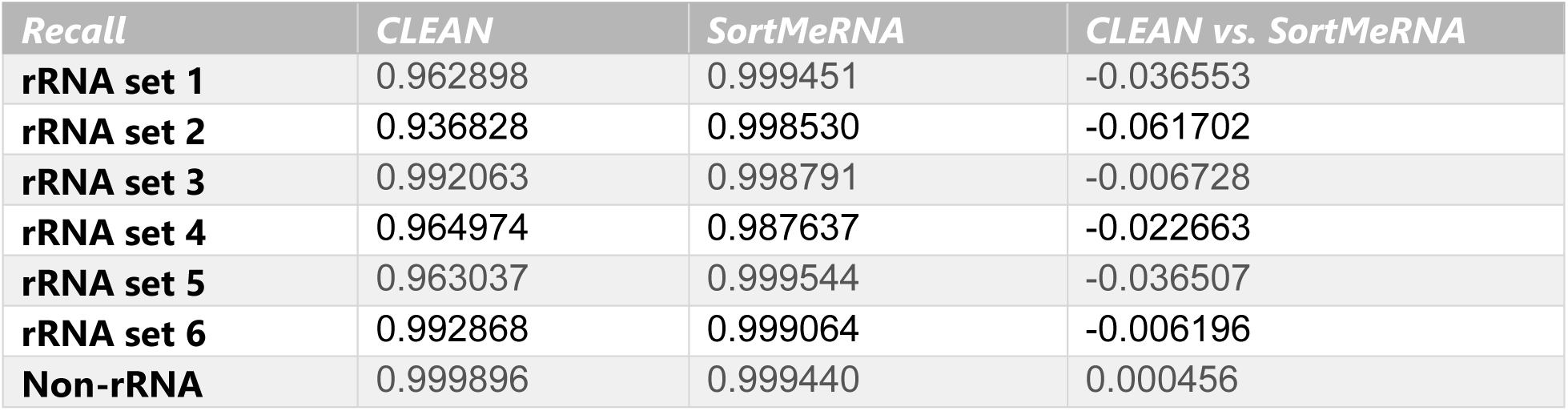
Recall comparison of CLEAN and SortMeRNA v4.3.4 for six simulated datasets consisting of 1 Mio Illumina rRNA reads each and one dataset consisting of 1 Mio Illumnia non-rRNA reads. CLEAN has a slightly decreased recall than SortMeRNA for the rRNA datasets; however, it is much faster (see **Supplementary Figure 4**).

**Supplementary Table 2.**
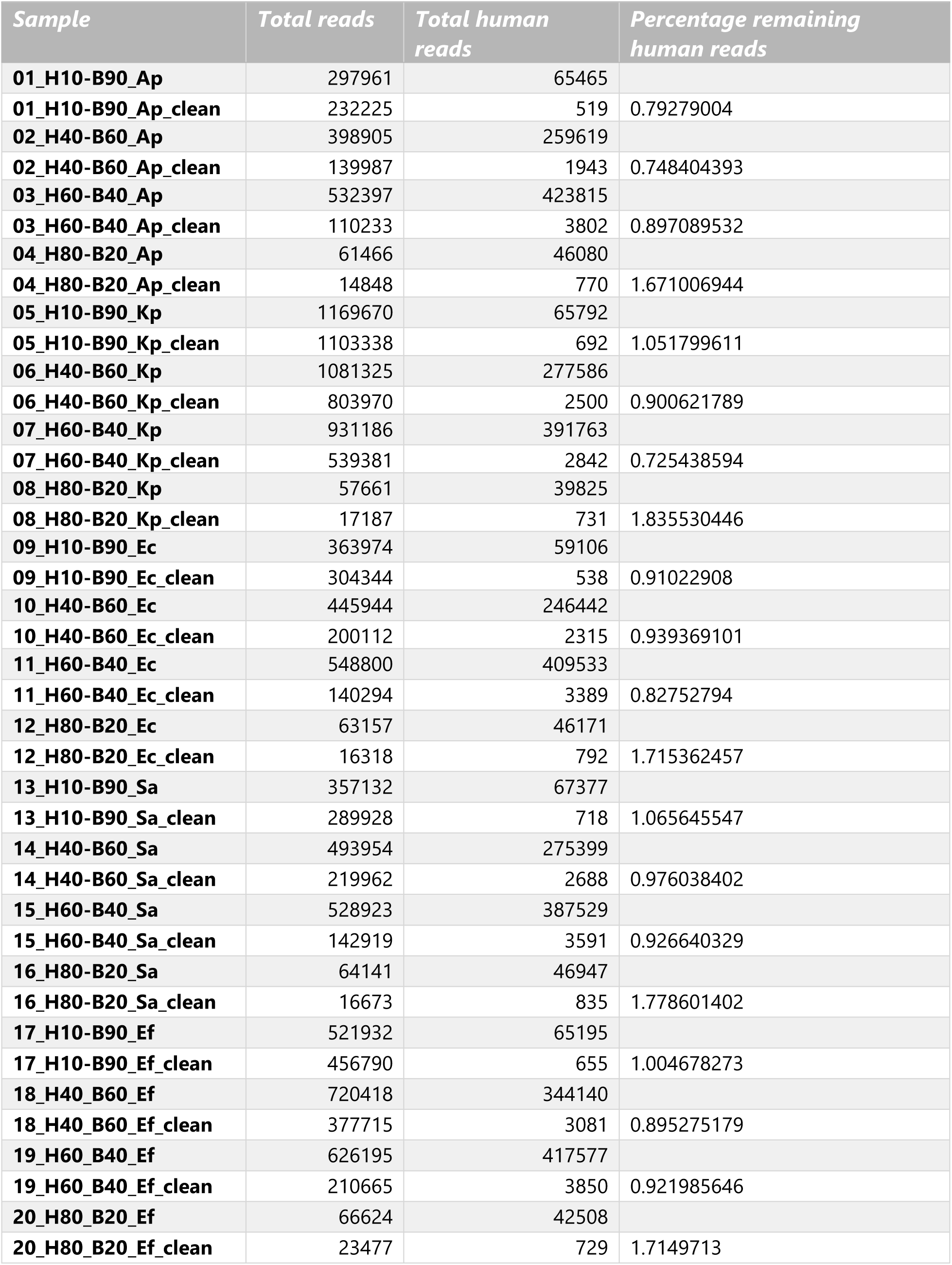
List of reads per sequencing run before and after CLEAN classified with Kraken 2. H — % of human DNA spike-in; B — % of bacteria DNA; Ap — *Acientobacter pittii*; Kp — *Klebsiella pneumoniae*; Ec — *Escherichia coli*; Sa — *Staphylococcus aureus*; Ef — *Enterococcus faecium*.

